# Long read sequencing reveals novel isoforms and insights into splicing regulation during cell state changes

**DOI:** 10.1101/2021.04.27.441628

**Authors:** David J Wright, Nicola Hall, Naomi Irish, Angela L Man, Will Glynn, Arne Mould, Alejandro De Los Angeles, Emily Angiolini, David Swarbreck, Karim Gharbi, Elizabeth M Tunbridge, Wilfried Haerty

## Abstract

Alternative splicing (AS) is a key mechanism underlying cellular differentiation and a driver of complexity in mammalian neuronal tissues. However, understanding of which isoforms are differentially used or expressed and how this affects cellular differentiation remains unclear. Long read sequencing allows full-length transcript recovery and quantification, enabling transcript-level analysis of AS processes and how these change with cell state. Here, we utilise Oxford Nanopore Technologies sequencing to produce a custom annotation of a well-studied human neuroblastoma cell line and to characterise isoform expression and usage across differentiation. We identify many previously unannotated features, including a novel transcript of the voltage-gated calcium channel subunit gene, *CACNA2D2*. We show differential expression and usage of transcripts during differentiation, and identify a putative molecular regulator underlying this state change. Our work highlights the potential of long read sequencing to uncover previously unknown transcript diversity and mechanisms influencing alternative splicing.

## INTRODUCTION

The complex suite of processes that occur during transcription gives rise to a staggering diversity of protein structures, molecular interactions and cell fates. Alternative splicing (AS) allows different transcripts to be generated from a single gene. Differential transcript expression (the overall abundance of a given transcript) or transcript usage (the abundance of a given transcript relative to that of others produced from the same gene) are key mechanisms for regulating cell lineage commitment and function (Breschi et al. 2020; Chepelev & Chen 2013). In vertebrates, AS is particularly prominent in the brain, and regulates multiple aspects of neurodevelopment including neurogenesis, synaptogenesis, cellular migration and axon guidance (Grabowski 2011; Ule et al. 2005; Raj & Blencowe 2015) in a temporally precise manner (Weyn-Vanhentenryck et al. 2018; Liu et al. 2018; Burke et al. 2020). These neurodevelopmental processes are defined by ordered switches in exon usage and expression across a spectrum of genes, controlled by a suite of highly specific RNA-binding proteins (RBPs) such as NOVA2 (Saito et al. 2019), PTBP1 and PTBP2 (Boutz et al. 2007; Linares et al. 2015; Keppetipola et al. 2012). A number of more ubiquitous RBPs may also help regulate these neuronal AS events, though which ones and what specific roles they play remain poorly understood (Jackson et al. 2020; Gallego-Paez et al. 2017).

Since AS can give rise to mRNAs that encode protein isoforms that exhibit distinct, or even opposing effects, it is essential to understand an individual gene’s products at transcript-level resolution (Clark et al. 2007; Yi et al. 2018; Liu et al. 2018; Yuste et al. 2020). However, the diversity of full-length transcripts remains poorly understood, as exemplified by the recent study of the L-type voltage gated calcium channel (VGCC) gene, *CACNA1C* (Clark et al. 2020). Furthermore, many unknowns remain as to the nature and regulation of changes in transcript expression during differentiation and development. For example, are there pronounced switches in primary transcript expression in a few key genes, or more nuanced expression differences across the transcriptome? Furthermore, although some of the molecular mechanisms that drive the observed ‘switches’ in transcriptional profiles occurring during lineage commitment have been identified, many of these processes remain to be determined.

As well as being of importance for understanding normal developmental processes, AS is also of clinical relevance, since aberrant transcriptional processes are implicated in many diseases (Scotti & Swanson 2016). Disease-associated mutations can directly affect AS by disrupting existing splice sites and/or forming novel or cryptic sites, as observed in the VGCC *CACNA1A* gene in Episodic Ataxia Type 2 (Jaudon et al. 2020). Alternatively, AS can alter disease presentation, as is seen in the case of Timothy Syndrome where the localisation of the disease-causing mutation in one of two mutually exclusive exons of *CACNA1C* determines syndrome severity (Splawski et al. 2004). Global changes in differential isoform expression are also associated with psychiatric conditions (Gandal et al. 2018).

Transcriptome profiling and annotation are essential first steps in investigating gene, isoform and exon expression or usage differences during cell differentiation. Until recently profiling was hampered by technological constraints, relying on short read sequencing technology (Stark et al. 2019; Wang et al. 2009). Whilst short read technologies provide cheap, accurate and high-coverage reads, with good differential expression analysis power (Wang et al. 2009), their ability to resolve and quantify fulllength transcripts is inherently limited (Byrne et al. 2017). In this context, the advent of long read technologies has rapidly improved our ability to characterise the transcriptome (Byrne et al. 2017, 2019) revealing, for example, the complexity of the transcriptional landscape of the mammalian brain (Wang et al. 2019; Sessegolo et al. 2019).

Here, we use long read sequencing to identify and quantify isoforms during a cellular state change; specifically, during the differentiation of the well-validated SH-SY 5Y neuroblastoma line into neuronlike cells. SH-SY5Y cells exhibit a stable genomic structure and have been widely used to investigate AS mechanisms and cellular differentiation from a neuroblast-like state (Kovalevich & Langford 2013; Shipley et al. 2016; Agholme et al. 2010; Truckenmiller et al. 2001), into a neuronal-like state (Forster et al. 2016; Mendsaikhan et al. 2018). We generated a custom high-coverage long read transcriptome annotation (using Oxford Nanopore Technology [ONT] cDNA sequencing), validated with orthogonal short read sequencing (Illumina paired-end short read) data. We identified novel transcriptomic features and performed differential expression and usage analyses to identify transcripts that show variation during differentiation, as well as identifying a novel putative molecular regulator underlying this state change.

## RESULTS AND DISCUSSION

### ONT reads accurately detect differential isoform expression

Using the Oxford Nanopore GridION platform, we generated on average 10,691,538 QC-passed reads per sample (± 1,751,518.6 SD). We also generated an average of 105,349,119 lllumina read pairs per sample (± 17,312,599.92 SD, Table S1). Whilst the utility of long read sequencing for recovering full length transcripts is widely accepted, there remains uncertainty as to the sensitivity of this technology for differential expression analysis. It is therefore important to assess the performance of long read vs short read sequencing in both transcript quantification and its application to differential expression studies (Sessegolo et al. 2019). We investigated the ability of the ONT data to detect Sequin spike-ins (Hardwick et al. 2016) of known concentration in a set of two different concentration mixes. We found that ONT limit of quantification (minimum transcript concentration) was 0.059 attomol/μl for mixA and 0.27 attomol/μl for mixB (Figs. 1A, 1B). By downsampling the short read data to the ONT average nucleotide coverage, we show there is similar power to detect transcripts by ONT and downsampled short read, although ONT scores lower than the complete short read data (Table S2). Next, we directly assessed the ability of the ONT reads to detect differential isoform expression. We calculated expected log2 fold-change (logFC) from the differences in concentrations between Sequins in mixA and mixB and compared this with observed logFC from our differential expression pipeline (see Methods). There was a strong correlation between expected and observed logFC of R^2^ = 0.973, p-value = 2.2e^−16^ (Fig. 1C), demonstrating that the ONT data can be used to detect differential expression over a broad range of logFC values. Collectively, our results indicate that our ONT data is sufficiently sensitive and powered to detect all but the lowest concentration transcripts and, further, that the ONT reads are suitable for differential isoform expression analysis. These results add to the growing literature highlighting the importance of assessing suitability of long read sequencing for differential expression.

**Fig 1.**
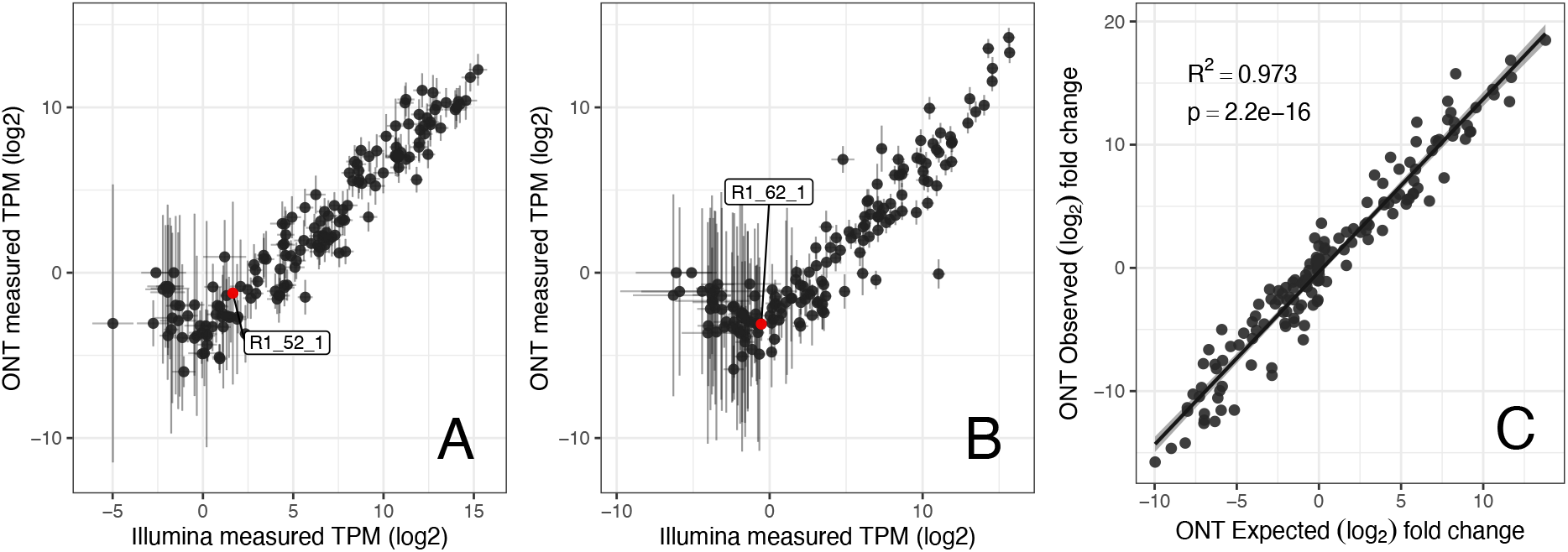
Comparison of Sequin spike-in sensitivity of detection between ONT (long read) and short read sequencing (Illumina paired-end, SRS) sequencing for Sequin synthetic spike-ins MixA (**plot A**) and MixB (**plot B**). Labelled sequin (red point) in each plot is at ONT limit of quantification (LOQ) in each mix; mix A 0.059 attomol/μl and mix B 0.27 attomol/μl. Plot C shows correlation of ONT differential isoform observed vs observed expression (log2 fold change +1) of synthetic Sequin spike-ins of known concentration. Pearson correlation coefficient is displayed along with a linear regression trend line with standard error in pale grey.

### TALON custom long read annotation reveals novel features of the human transcriptome

The TALON custom annotation provided a total of 3,274 novel transcripts prior to validation using short read sequencing. We found short read support for 2,567 of the 3,274 (78.41%) novel transcripts recovered from the ONT read data (Fig. 2) by stringent removal of transcripts that contained a novel exon lacking at least 15 reads depth across 75% of its length (see methods). The supported novel transcripts collectively include a total of 49 novel cassette exons (18 frame-conserving) along with 928 and 1046 novel 5’ and 3’ splice sites respectively, with 464 instances of exons exhibiting both a novel 5’ and 3’ splice site. Additionally, we identified 92 novel junctions between previously annotated splice sites. In total 929 (36.19%) of the validated novel transcripts were putatively coding; either frame-conserving, or assumed to be coding via CPAT (Wang et al. 2013) assessment, whilst 1638 (63.81%) were assumed noncoding due to either induction of a frameshift, a noncoding parent gene or via noncoding classification from CPAT (fig. 2 for full breakdown).

**Fig 2.**
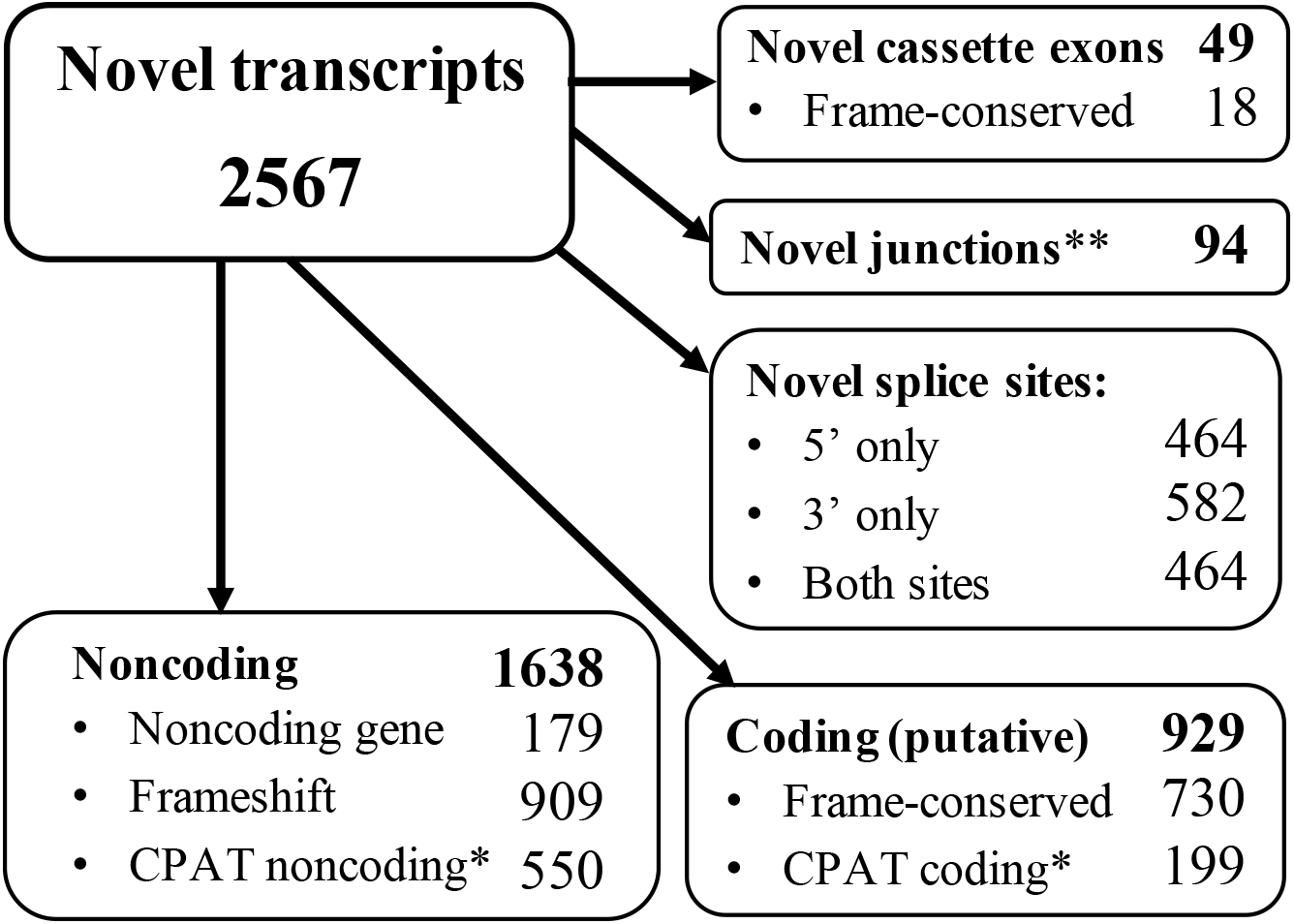
Breakdown of novel transcripts identified using ONT long reads and TALON custom transcriptome annotation. Cassette exons are previously unannotated positions. *CPAT assessment of CDS coding probability (CP ≥ 0.364). **Novel junctions are previously unannotated junctions identified between existing exonic parts. All novel assessments are relative to Gencode v29 human transcriptome annotation.

Our stringent filtering criteria and validation likely results in an underestimate of the true quantity of novel features present. Despite this, our long read sequencing approach still identifies >3000 novel transcripts, nearly a thousand of which are putatively coding. Collectively, our data highlight the extent of previously undescribed transcriptome diversity, even within a highly specialised (and well-studied) cell model. Our work concurs with the growing body of other studies using long reads for transcriptome assessment (Gleeson et al. 2020; Sessegolo et al. 2019, Soneson et al. 2019); relying on short reads substantially underestimates transcriptome diversity.

### ONT differential gene expression supports neuron-like characteristics of differentiated SH-SY5Y cells

Differential gene expression analysis revealed 4,239 genes differentially expressed (FDR q < 0.05) between differentiation states, with 2,041 and 2,198 genes overexpressed in undifferentiated and differentiated cells, respectively (Table 1, Fig 3A). We performed FUMA analyses to explore the functional significance of these genes. The upregulated genes in differentiated cells showed greatest overlap with those upregulated in brain compared with other tissue types (padj= 3.4 x 10^−41^), whilst for those more highly expressed in undifferentiated cells, there was overlap with those downregulated in brain (padj = 2.8 x 10^−45^, second only to pancreas: padj = 1.8 x 10^−45^). Gene Ontogeny biological pathway terms showing differential expression across differentiation included neurogenesis, neuron development, cell differentiation and regulation of nervous system development (Table S5).

**Fig 3.**
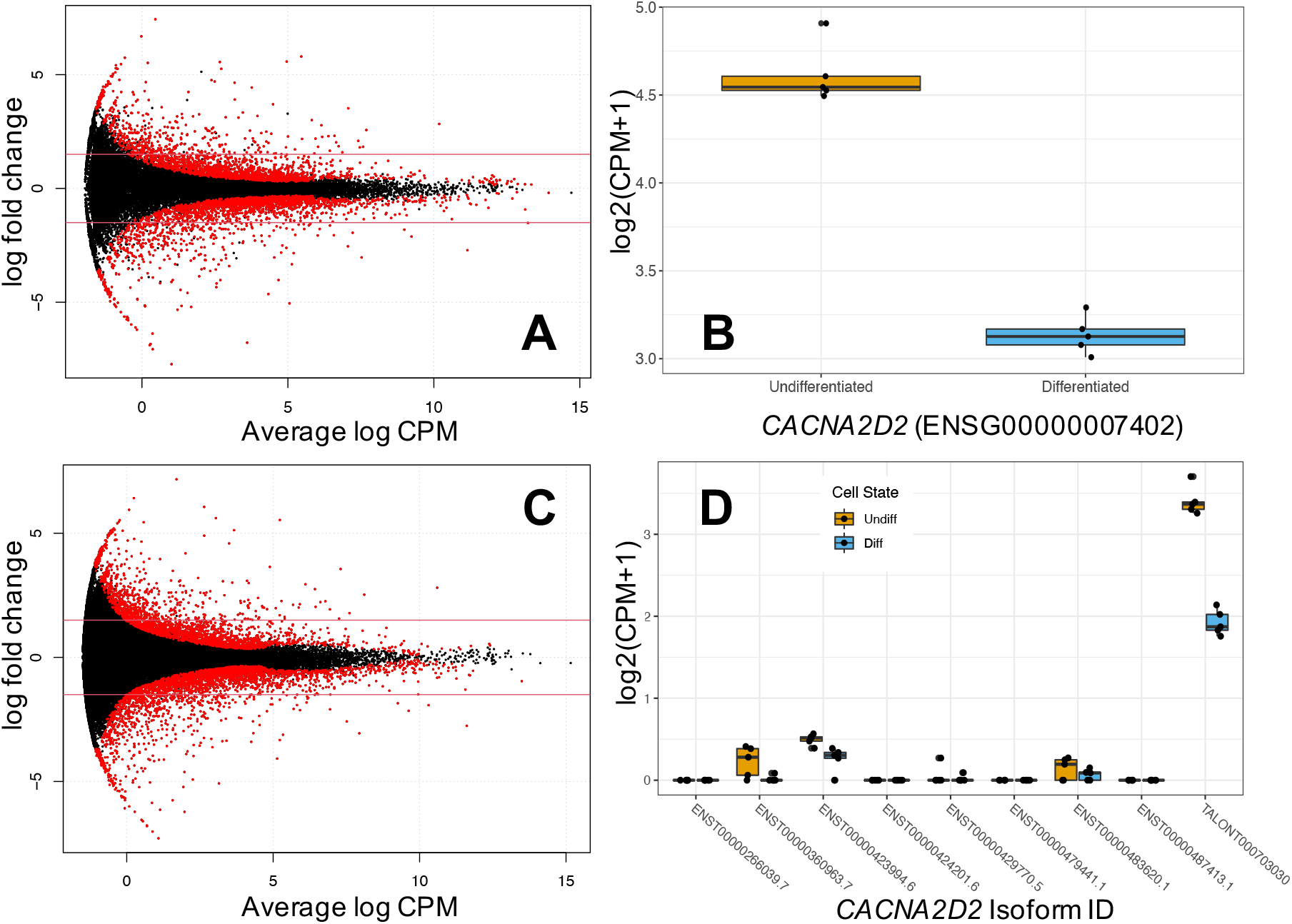
Panel of **A)** gene-level differential expression (DE) smear plot, solid red lines highlighting ± 1.5 logFC threshold, **B)** gene-level DE of *CACNA2D2* during differentiation, **C)** isoform-level DE smear plot with ± 1.5 logFC threshold, **D)** *CACNA2D2* isoform expression, showing novel TALON isoform with highest read count. Red points on smear plots indicate significant differential expression (FDR < 0.05). Boxplots display median and IQR.

**Table 1.** Showing number of genes and transcripts differentially expressed between undifferentiated and differentiated cells, after multiple testing correction using ONT long read counts (FDR < 0.05). Bracketed numbers refer to the portion of total that are TALON-identified novel transcripts. U = undifferentiated and D = differentiated cells, with arrows displaying expression directionality.

| Metric | Count |  |
| --- | --- | --- |
|  | Gene level | Transcript level (Talon) |
| Total features assessed | 32977 | 99067 (1855) |
| Differentially Expressed | 4239 | 5456 (197) |
| ↑U ↓D (all > 0 log<sub>2</sub>FC) | 2041 | 2390 (67) |
| ↑D ↓U (all < 0 log<sub>2</sub>FC) | 2198 | 3066 (130) |

### Isoform-level differential expression reveals novel diversity

At the isoform-level, we detected differential expression of 5,456 transcripts (FDR q-value < 0.05) in 4,416 genes (Table 1, Fig 3C) across cell state. It is important to note that, of the 4,416 genes including differentially expressed isoforms, 1,276 were not differentially expressed at the gene level. Analysing expression at the transcript level therefore provides a more accurate overview of transcriptional dynamics during cellular differentiation. FUMA analysis of the genes that show differential transcript expression (DTE) in the absence of differential gene expression (DGE) identified intracellular trafficking and macromolecule localisation, and post-transcription and -translation processing mechanisms. These findings highlight the importance of post-transcriptional mechanisms in cellular differentiation and development and emphasise the importance of understanding changes at the transcript level, as key processes may be obscured at the gene level.

### Long read transcriptome annotation reveals a novel CACNA2D2 isoform and exon

As a specific example of the power of long read sequencing for uncovering isoform diversity, we identified a novel transcript (TALONT000703030) of the VGCC α_2_δ_2_ subunit gene, *CACNA2D2*, which is truncated (18,731bp), compared to most previously annotated isoforms (87,831bp - 141,442bp, but note ENST00000483620 at 2927bp, Fig. S2). This transcript is designated novel as it includes a novel first exon (chr3:50381499-50381529), with a potential in-frame start site (chr3:50381507-50381509) leading to an open reading frame. Orthogonal short read validation showed that this is not an artefact of ONT sequencing (Fig. S3). Evidence for the existence of the exon can also be found in human cortex GTEx v.8 (Carithers et al. 2015) RNA-seq data (N=27 samples assessed from N=21 individuals, Fig. S4) suggesting it is not unique to SH-SY5Y cells. Surprisingly, TALONT000703030 was the most highly expressed *CACNA2D2* isoform in the undifferentiated state and was downregulated in differentiated cells (Fig. 3D & isoform differential expression results), consistent with an overall downregulation of *CACNA2D2* at the gene level following differentiation (Fig. 3B & gene differential expression results). The *CACNA2D2* encoding subunit is vital to normal channel trafficking and has complex regulatory effects on VGCC currents (Dolphin 2016; Gao et al. 2000). Mutations in *CACNA2D2* have long been studied for links with epileptic and cerebellar ataxic phenotypes (Brill et al. 2004; Barclay et al. 2001), and previously described truncated proteins of *CACNA2D2* resulting from mutations exhibit abnormal function (Brodbeck et al. 2002). It remains to be determined whether the novel isoform presented here is functional and, if so, how its function differs from annotated isoforms. A CPAT (Wang et al. 2013) analysis reveals it to be potentially coding and initial Phyre2 (Kelley et al. 2015) predictions suggest it would result in a cleaved protein structure (Fig. S5). Alternative splicing of other VGCC subunits is a key means of generating functionally distinct channels (Hofmann et al. 2014). However, the novel isoform lacks the signal peptide (SignalP 5.0, Armenteros et al. 2019) required for membrane insertion, and therefore is unlikely to encode a functional VGCC α_2_δ subunit (Dolphin 2018). Given that this transcript is downregulated as cells differentiate into neuron-like cells, it is possible that the switch from a non-functional to a functional α2δ subunit represents a means of regulating VGCC signalling, potentially preventing its inadvertent activation in undifferentiated cells.

### DTU analysis reveals splicing regulators relevant to SH-SY5Y differentiation

Whilst isoform-level expression aids the identification of important proteins and functional changes during differentiation, understanding differential transcript usage (i.e. the abundance of a given transcript relative to that of others produced from the same gene) may provide further insights. Using IsoformSwitchAnalyzeR v.1.11.3 (Vitting-Seerup & Sandelin 2019) we identified 104 isoform switches (each affecting a single gene) considered to have putative functional consequences (see methods) across SH-SY5Y differentiation (Table S4). The implicated genes include the histone deacetylase gene, *HDAC4*, and *RACGAP1*, which encodes Rac GTPase Activating Protein 1, a protein critical for axon morphogenesis (Warga et al. 2016).

We identified RNA binding motif protein 5 (*RBM5*), a ubiquitous splicing regulator (Jackson & Kochanek 2020), among the genes showing differential transcript usage. Upon differentiation, we observed a switch from a noncoding to a coding transcript for this splicing factor (Fig. 4). Based on this observation, we intersected the DTU genes against a list of human RBPs with eCLIP-seq binding data from a HepG2 line generated by the ENCODE project (Van Nostrand et al. 2020) to identify putative *RBM5* targets. A hypergeometric test revealed enrichment for binding targets of *RBM5* in our 104 DTUs (fold enrichment = 2.17, p-value = 5.87×10^−14^). This suggests that RBM5 may play a role in splicing regulation during differentiation of SH-SY5Y cells.

**Fig 4.**
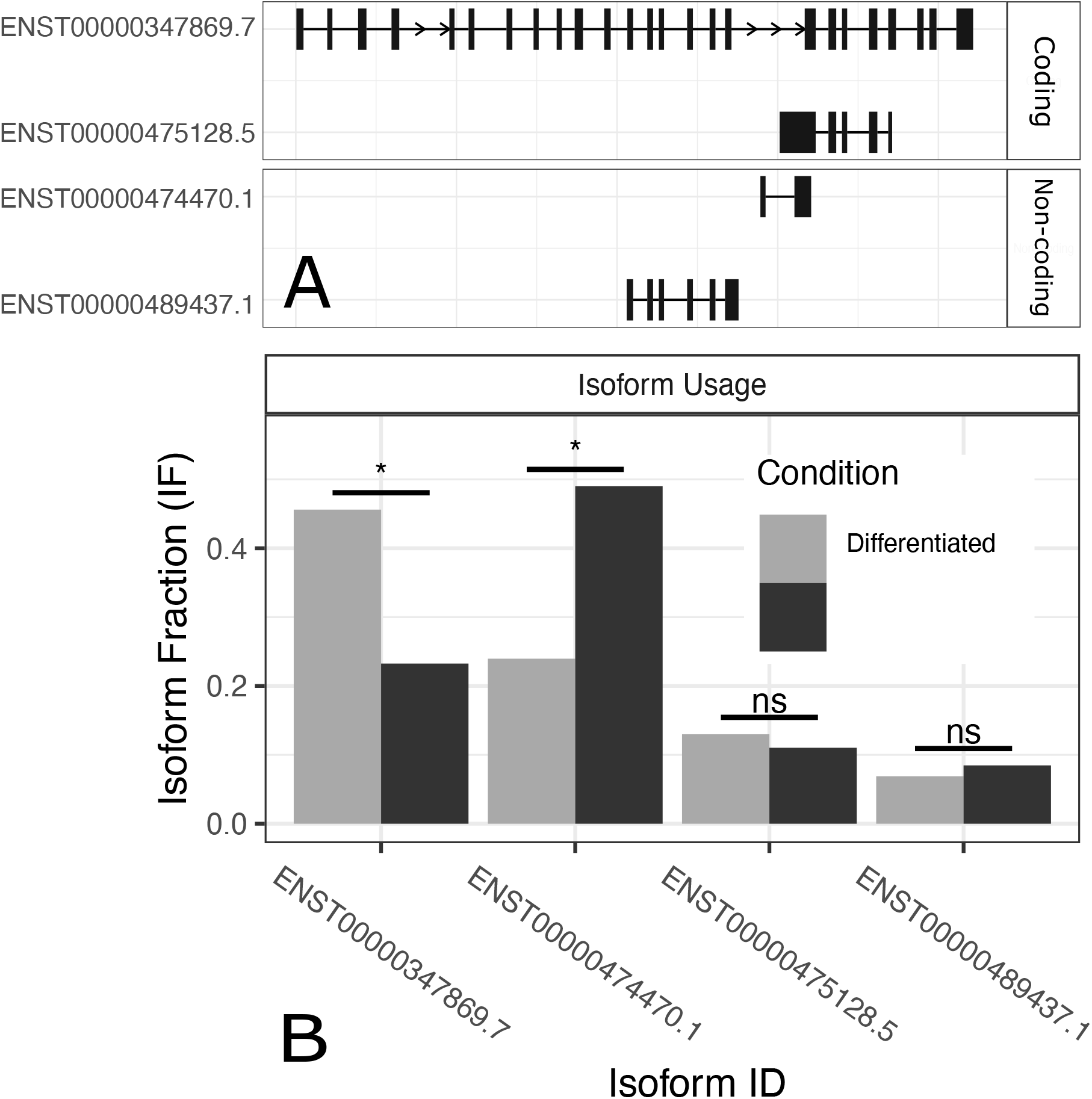
Panel of **A)** known *RBM5* isoforms with relative size and exon composition and **B)** differential transcript usage (DTU) of known *RBM5* isoforms across differentiation displaying a switch from a truncated, non-coding *RBM5* isoform ENST00000474470 to full-length coding isoform ENST00000347869. Figures modified from IsoformSwitchAnalyzeR (Vitting-Seerup, 2017) output.

The function of *RBM5* in neurogenesis is poorly understood (Jackson & Kochanek 2020), although its roles in apoptosis and tumour suppression (Xu et al. 2019; Zhang et al. 2019; Sutherland et al. 2010; Bechara et al. 2013) and spermatogenesis (O’Bryan et al. 2013) are well documented. We show a clear switch from a truncated, non-coding *RBM5* isoform ENST00000474470 to the full-length coding isoform ENST00000347869 during differentiation (Fig. 4A, 4B). Our hypergeometric enrichment test revealed that 69 of the 104 DTU genes are targeted by *RBM5* binding in the ENCODE data. Future work needs to further investigate how changes in *RBM5* regulation impacts isoform expression through experimental confirmation of *RBM5* binding targets in SH-SY5Y cells, and mutagenesis of *RBM5* and its binding sites. Thus, by integrating novel annotations and isoform expression dynamics using long read RNA sequencing with existing (and emerging) epigenomic data sets (RBP e-CLIP data, in this case), we identified candidates in the regulatory network involved in SH-SY5Y differentiation.

Together, our findings demonstrate the power of long read sequencing for the characterisation and quantification of genes at the isoform level. We demonstrate that current annotations remain far from complete, even in the well-studied SH-SY5Y cell line. We show the utility of this model system and our technical approach for studying fundamental molecular processes underlying changes in cell state. Finally, our findings indicate that changes in *RBM5* expression profiles may act as a molecular mechanism for the coordination of these changes, paving the way for future functional studies.

## MATERIALS AND METHODS

### Sampling and Sequencing

#### Cell culture and neuronal differentiation

A total of 10 technical replicates of human neuroblastoma SH-SY5Y cells were cultured in neurobasal media (Gibco 21103-049) supplemented with B-27 Plus (ThermoFisher). Retinoic acid (SigmaAldrich) was added to five replicates to a final concentration of 10mM, to induce cell differentiation to a neuronlike state; whilst five replicates were cultured to confluence in standard media. Cells were washed with phosphate buffered saline and harvested in QIAzol (Qiagen) to preserve RNA, before being stored at −80°C until RNA extraction.

#### RNA extraction and spike-in control

Total RNA was purified from the 10 replicate cell cultures using a Direct-zol RNA Miniprep Plus kit (Zymo Research), according to the manufacturer’s instructions. Internal controls were employed to assess long read sequencing sensitivity to transcript detection and differential expression. Sequin synthetic spike-ins (Hardwick et al. 2016) are designed in a range of sizes and utilised in two separate mixes (MixA v.2 & MixB v.1) of known, contrasting concentrations (see data repository for details). These were spiked into the 10 replicates in an alternating fashion, so that MixA and MixB were represented in both cell states to enable internal control. The Sequins were spiked in at 1% of the total RNA input amount (https://www.sequinstandards.com/). These were added to the native RNA prior to reverse transcription, as per manufacturer’s instructions.

#### cDNA long read sequencing (ONT)

200ng total RNA per sample was processed using the cDNA-PCR Sequencing Kit (Oxford Nanopore Technologies, SQK-PCS109). The first-strand reaction was prepared according to the ONT cDNA-PCR Sequencing protocol provided by the manufacturers (SQK-PCS109, https://community.nanoporetech.com/protocols/cdna-pcr-sequencing_sqk-pcs109/v/PCS_9085_v109_revJ_14Aug2019?devices=gridion). Post first-strand, the reaction was snap cooled and then incubated to 42°C while 8μl of second-strand switching primer (SSP) master mix was added (prepared according to the protocol). This was mixed and incubated at 42°C for 2 minutes. 1μl of Maxima H reverse transcriptase was then added to the second-strand reaction, maintained at 42°C. The whole second-strand reaction was then mixed and incubated at 42°C for 90 minutes. The reaction was then inactivated at 85°C for 5 minutes and held at 4°C until ready to proceed to PCR. After reverse transcription, the reaction was split into four aliquots in preparation for four replicate PCR tubes to generate sufficient products while minimising the risk of over-amplification. The reagents for PCR were prepared according to the cDNA-PCR Sequencing (SQK-PCS109) protocol and incubated under the following conditions; initial denaturation at 95°C for 30 sec followed by 15 cycles of: denaturation at 95°C for 15 sec, annealing at 62°C for 15 sec, extension at 65°C for 5 mins; with a final extension of 65°C for 6 mins and hold at 4°C. Post-PCR each PCR aliquot was exonuclease I (NEB) treated. Aliquots were then pooled per sample and bead concentrated (Beckman Coulter, AMPure XP, A63880) prior to sequencing. The cDNA was quantified using High Sensitivity Qubit assays (ThermoFisher, Q32854) and sized using the 2100 Bioanalyzer instrument (Agilent Technologies, cat. no. G2939BA) High Sensitivity DNA assay (Agilent, 5067-4626). 250ng cDNA per sample was prepared for rapid adapter ligation (approximately 190 fmol per sample).

MinION flowcells underwent QC prior to library construction. The PCR cDNA libraries were prepared for sequencing and the respective MinION flowcells were primed following the cDNA-PCR Sequencing (SQK-PCS109) protocol using the PromethION flow cell priming kit (Oxford Nanopore Technologies, EXP-FLP001.PRO.6). Five PCR-cDNA libraries were sequenced in parallel per run of the GridION instrument with GridION software v.18.12.4 (Oxford Nanopore Technologies), using one Flowcell per library. Basecalling for all PCR-cDNA libraries was completed in real-time using MinKNOW on the GridION and maximum data acquisition time of 48 hours was conducted for all ten flowcells. ONT fast5 files were first converted to fastq using guppy v.3.2.2 (https://community.nanoporetech.com) before QC using MultiQC (Ewels et al. 2016).

#### cDNA short read sequencing

Illumina library preparation and sequencing were carried out by the Genomics Pipelines team at Earlham Institute. 1μg of total RNA per sample was processed using the NEBNext Ultra II Directional RNA library prep kit from NEB (E7760L) utilising the NEBNext Poly(A) mRNA Magnetic Isolation Module (E7490L) with NEBNext Multiplex Oligos for Illumina® (96 Unique Dual Index Primer Pairs, E6440S) at a concentration of 10μM. The RNA was purified to extract mRNA with a Poly(A) mRNA Magnetic Isolation Module. Isolated mRNA was then fragmented and cDNA was synthesised for the first strand. The second strand synthesis process removes the RNA template and synthesises a replacement strand to generate ds cDNA. Directionality is retained by adding dUTP during the second strand synthesis step and subsequent cleavage of the uridine-containing strand using USER Enzyme (a combination of UDG and Endo VIII). NEBNext Adaptors were ligated to end-repaired, dA-tailed DNA. The ligated products were subjected to a bead-based purification using Beckman Coulter AMPure XP beads (A63880). Adaptor ligated DNA was then enriched by receiving 10 cycles of PCR; 30 secs at 98°C, 10 cycles of: 10 secs at 98°C, 75 secs at 65°C, 5 mins at 65°C, final hold at 4°C. Barcodes were incorporated during the PCR step. The resulting libraries underwent QC using PerkinElmer GX and a High Sensitivity DNA chip (5067-4626), the concentration was determined with a High Sensitivity Qubit assay (Q32854) or plate reader. The final libraries were pooled, a qPCR was performed using a KAPA Illumina ABI library quantification kit (Roche Diagnostics, 7960204001) on a StepOne q-PCR machine (ThermoFisher), and then these were prepared for sequencing.

The library pool was diluted to 0.65nM with 10mM Tris (pH 8.0) in a volume of 18μl before spiking in 1% Illumina PhiX Control v.3. This was denatured by adding 4μl 0.2N NaOH and incubating at room temperature for 8 mins, after which it was neutralised by adding 5μl 400mM Tris (pH 8.0). A master mix of EPX1, EPX2, and EPX3 from Illumina’s Xp 2-lane kit was made and 63μl added to the denatured pool leaving 90μl at a concentration of 130 pM. This was loaded onto a NovaSeq S1 flow cell using the NovaSeq Xp Flow Cell Dock. The flow cell was then loaded onto the NovaSeq 6000 along with a NovaSeq 6000 S1 cluster cartridge, buffer cartridge, and 200 cycle SBS cartridge. The NovaSeq had NVCS v.1.6.0 and RTA v.3.4.4 and was set up to sequence 100bp PE reads. The data were demultiplexed and converted to fastq using bcl2fastq2 v.2.20 (Illumina). Illumina data underwent QC with MultiQC v.1.5 (Ewels et al. 2016) and adaptors were removed with trim galore v.0.5.0 (Krueger 2015) with default parameters.

#### Custom transcriptome annotation and validation

The cDNA ONT reads were aligned to the human genome (hg38, modified to include an artificial chromosome containing the sequins spike-ins as contiguous transcripts) using minimap2 v.2.17 (Li 2018) in sam MD-tag aware mode. Only primary alignments were retained. We then employed TALON v.5.0 (Wyman et al. 2020), a technology-agnostic pipeline that leverages long reads to build a custom transcriptome annotation. By exploiting the ability of long reads to detect full-length transcripts, TALON identifies novel features through comparison to an existing reference annotation. First, the sam files were passed to TranscriptClean v.2.0.2 (Wyman & Mortazavi 2019) for correction of read microindels (<5bp) and mismatches, though any non-canonical splice junctions were retained for downstream analyses, as novel features can often be found with non-canonical junctions (Sibley et al. 2016). The reads were checked for internal priming artifacts, a known issue with oligo-dT poly(A) selection methods (Roy & Chanfreau 2020), using a T-window size of 20bp (equivalent to the primer T sequence) and removed. Read annotation was performed with TALON, using the human Gencode v.29 reference annotation gtf with minimum alignment identity = 0.9 and coverage = 0.8. All 10 cDNA ONT replicates were utilised to maximise recovery of novel features. Identified transcripts were subsequently filtered using a minimum count threshold of N=5 reads in K=3 samples. As we expect to find lowly expressed isoforms in both biological conditions (N = 5 replicates per condition), K=3 was selected to balance sensitivity with accuracy. An updated annotation was produced using this filtered set of transcripts. The TALON custom gtf contains only features detected with reads present in the dataset, so a complete custom transcriptome annotation was compiled by merging the reference and TALON gtfs.

We then employed a series of quality control and validation steps. Novel antisense transcripts perfectly matching existing gene models were removed. Novel features were then validated by assessing short read exon coverage with bedtools v.2.28.0 (Quinlan & Hall 2010) using the orthogonal cDNA short read data genome-aligned with HISAT v.2.0.5 (Kim et al. 2015) and a minimum threshold of 15 reads depth over at least 75% of exon length. Any entries containing novel exons that failed this validation were removed from the gtf. The resulting validated annotation file is herein referred to as the TALON gtf (fig. S1 for full pipeline). Novel features were explored utilising a series of custom scripts to assess frameshifting and coding potential with CPAT v.2.0 (Wang et al. 2013) and publicly available data within the UCSC browser and associated databases (Kent et al. 2002). The *CACNA2D2* novel exon was validated by orthogonal short read coverage of all 10 samples using bedtools coverage. Evidence was additionally sought in human cortex RNA-seq data by accessing 27 accessions from 21 individuals in the publicly available GTEx database in the same manner.

### Differential expression analyses

#### Sequin spike-in detection & ONT DE sensitivity

Sensitivity in detecting isoform DE using ONT was assessed by a) finding the threshold of detection for each Sequin mix, b) comparing observed vs expected logFC and c) comparing with short read data. Read TPM (transcripts per million) was calculated with Salmon v.0.13.1 (Patro et al. 2017) and analysed with Anaquin v.2.8.0 (Wong et al. 2017). Anaquin finds the limit of quantification (LOQ) concentration for each Sequin mix by fitting linear regression on the entire dataset, minimising the total sum of squares of differences between variables (Wong et al. 2017). This was also performed for both the full short read data and a version downsampled to equivalent ONT average nucleotide coverage using bedtools. Differential expression analyses were performed with edgeR v.3.30.3 (McCarthy et al. 2012) in R v.4.0.2 (R Core Team 2016). Counts (numReads) were generated from unaligned reads at transcript-level using Salmon in mapping-based mode for the Sequins in each replicate. We then utilised a standard differential expression pipeline (detailed below). The differential expression regression model was specified by splitting the data into Sequin MixA and MixB accordingly. Expected log-fold change (logFC) was calculated as log2(MixB/MixA-1)+1 for each Sequin spike-in, for direct comparison with the observed logFC calculated with edgeR.

#### Gene and transcript-level differential expression

Differential expression analyses were carried out at both transcript (DTE) and gene (DGE) level. ONT reads were mapped to the TALON transcriptome using minimap2 and quantified with Salmon in alignment-based mode and using 100 bootstrap replicates. Any novel transcripts from the Talon custom annotation that were not located on assembled chromosomes were removed to reduce any impact from counting errors associated with scaffold-only/duplicated transcripts. Transcript-level counts were then obtained by importing Salmon results with the EdgeR function catchSalmon(), using the bootstrap replicates to calculate and apply an overdispersion correction for each count. As ONT reads achieve relatively low coverage compared with short reads, any transcripts unexpressed/undetected across all the 10 samples were removed from the data but all other counts were retained. Gene-level counts were generated by subsequently removing these unexpressed/undetected transcripts from the Salmon quantification (quant.sf) files and importing these pre-filtered data directly to EdgeR with tximport v.1.16.1 (Soneson et al. 2015) and an isoform-to-gene conversion matrix built from the Talon gtf. We then utilised a standard differential expression pipeline in edgeR for both DTE and DGE. The data were TMM-normalised and the biological coefficient of variation (BCV) and multidimensional scaling (MDS) were both manually assessed for outliers and confounding variation in the dataset. In each analysis, a model matrix was specified and applied to a glm, with false discovery rate (Benjamini-Hochberg FDR) correction for multiple testing of the results. Both DTE and DGE results were also filtered at a threshold of ± 1.5 logFC (log2 fold change) and FDR < 0.05, to obtain the most differentially expressed subset of features in each case for downstream analyses.

#### Differential usage analyses

Differential transcript usage (DTU) was assessed using the R package IsoformSwitchAnalyzeR v.1.11.3 (Vitting-Seerup & Sandelin 2019) on the same transcript quantification input used for DTE and DGE. TPM abundances were imported using the scaledTPM function in tximport and imported into IsoformSwitchAnalyzeR. The DTU analysis was run in two parts; first non-expressed isoforms were removed, and switches calculated for each gene using DEXseq (Anders et al. 2012) and nucleotide and peptide outputs for each gene were created for protein assessment. Transcripts were assessed for coding potential with CPAT (Wang et al. 2013), protein domain assignment with PFam (Punta et al. 2012), signal peptide prediction with SingalP v.5.0 (Armenteros et al. 2019) and intrinsically disordered regions and binding regions with IUPred2A (Mészáros et al. 2018), using default parameters according to the IsoformSwitchAnalyzeR workflow. The second part of the IsoformSwitchAnalyzeR DTU analysis then leveraged these data to identify isoforms switches with potential functional consequences and provide visualisation using default functions.

#### Hypergeometric enrichment tests

The set of putative functionally consequential DTUs was checked for RBPs by intersection with a set of known RBPs assayed as part of the ENCODE project (Van Nostrand et al. 2020), revealing the presence of *RBM5*. Corresponding narrow-peak eCLIP bed data for *RBM5* were accessed using the ENCODE portal (Davis et al. 2018) for both HepG2 isogenic replicates in the ENCODE repository (*RBM5* accessions ENCFF176RGG and ENCFF998ACW downloaded 29/09/2020) and intersected using bedtools to find the most supported subset of binding targets. The intersection for both significant DTUs (N=104) and for total genes assessed (N=32325) with the eCLIP binding targets were then obtained by intersection with this subsetted list of ENCODE targets. A hypergeometric test for enrichment was performed using the phyper functionality in the R core package ‘stats’ v.4.0.2 (R Core Team 2016).

#### Ontology and functional association

To interpret the differentially expressed or used gene sets, we assessed gene ontology and known associations with neurologically relevant biology. We used the GENE2FUNC function in FUMA (Watanabe et al. 2017) to annotate the gene sets within a biological context. For transcripts, the corresponding Ensembl gene ID was used. In each case, the default thresholds of significance and ontology enrichment were applied. Analyses focused on tissue specificity analyses in GTEx v.8 30 tissue types and Gene Ontogeny (GO) Biological Processes.

## ACKNOWLEDGEMENTS & FUNDING

The work was funded by a UK Medical Research Council [MR/P026028/1] award to EMT and WH, supported by the Oxford Health NIHR Biomedical Research Centre. The author(s) acknowledge the support of the Biotechnology and Biological Sciences Research Council (BBSRC), part of UK Research and Innovation; this research was funded by the BBSRC Core Strategic Programme Grant (Genomes to Food Security) BB/CSP1720/1 and its constituent work packages(s) - BBS/E/T/000PR9818 (WP1 Signatures of Domestication and Adaptation) and Core Capability Grant BB/CCG1720/1 and the National Capability BBS/E/T/000PR9816 (Supporting EI’s ISPs and the UK Community with Genomics and Single Cell Analysis) and BBS/E/T/000PR9811 (NC4 Enabling and Advancing Life Scientists in data-driven research through Advanced Genomics and Computational Training). The authors would like to thank Norwich Research Park CiS team for technical assistance. This study makes grateful use of ENCODE Consortium data generated by the Yeo Lab UCSD and Genotype-Tissue Expression (GTEx) Project v.8 which is supported by the Common Fund of the Office of the Director of the National Institutes of Health, and by NCI, NHGRI, NHLBI, NIDA, NIMH, and NINDS. The views expressed are those of the authors and not necessarily those of the National Health Service, NIHR or the Department of Health.

## DATA AVAILABILITY

Short read (PRJEB44501) and long read (PRJEB44502) RNAseq fastq files were archived in ENA. Data files and analysis scripts available at: https://github.com/TGAC/SHSY5Y_differentiation or the corresponding author.

## AUTHOR CONTRIBUTIONS

WH, KG & EMT project lead and design, DW analysis and writing lead, KG sequencing lead, NI & WG sample preparation and sequencing, NH & AM cell culture, ALM, AA, & DS analysis. EA project support and management of WG. All authors contributed to and agreed to manuscript submission.

## Notes

### Competing Interest Statement

The authors have declared no competing interest.

## CITATIONS

Agholme L, Lindström T, Kågedal K, Marcusson J, Hallbeck M. 2010. An in vitro model for neuroscience: differentiation of SH-SY5Y cells into cells with morphological and biochemical characteristics of mature neurons. J. Alzheimers. Dis. 20:1069–1082.

Anders S, Pyl PT, Huber W. 2015. HTSeq--a Python framework to work with high-throughput sequencing data. Bioinformatics. 31:166–169.

Anders S, Reyes A, Huber W. 2012. Detecting differential usage of exons from RNA-seq data. Genome Res. 22:2008–2017.

Armenteros JJA et al. 2019. SignalP 5.0 improves signal peptide predictions using deep neural networks. Nat. Biotechnol. 37:420–423.

Barclay J et al. 2001. Ducky mouse phenotype of epilepsy and ataxia is associated with mutations in the Cacna2d2 gene and decreased calcium channel current in cerebellar Purkinje cells. J. Neurosci. 21:6095–6104.

Bechara EG, Sebestyén E, Bernardis I, Eyras E, Valcárcel J. 2013. RBM5, 6, and 10 differentially regulate NUMB alternative splicing to control cancer cell proliferation. Mol. Cell. 52:720–733.

Boutz PL et al. 2007. A post-transcriptional regulatory switch in polypyrimidine tract-binding proteins reprograms alternative splicing in developing neurons. Genes Dev. 21:1636–1652.

Breschi A et al. 2020. A limited set of transcriptional programs define major cell types. Genome Res. 30:1047–1059.

Brill J et al. 2004. entla, a novel epileptic and ataxic Cacna2d2 mutant of the mouse. J. Biol. Chem. 279:7322–7330.

Brodbeck J et al. 2002. The ducky mutation in Cacna2d2 results in altered Purkinje cell morphology and is associated with the expression of a truncated alpha 2 delta-2 protein with abnormal function. J. Biol. Chem. 277:7684–7693.

Burke EE et al. 2020. Dissecting transcriptomic signatures of neuronal differentiation and maturation using iPSCs. Nat. Commun. 11:462.

Byrne A et al. 2017. Nanopore long-read RNAseq reveals widespread transcriptional variation among the surface receptors of individual B cells. Nat. Commun. 8:16027.

Byrne A, Cole C, Volden R, Vollmers C. 2019. Realizing the potential of full-length transcriptome sequencing. Philos. Trans. R. Soc. Lond. B Biol. Sci. 374:20190097.

Carithers LJ et al. 2015. A Novel Approach to High-Quality Postmortem Tissue Procurement: The GTEx Project. Biopreserv. Biobank. 13:311–319.

Chepelev I, Chen X. 2013. Alternative splicing switching in stem cell lineages. Front. Biol. 8:50–59.

Clark MB et al. 2020. Long-read sequencing reveals the complex splicing profile of the psychiatric risk gene CACNA1C in human brain. Mol. Psychiatry. 25:37–47.

Clark TA et al. 2007. Discovery of tissue-specific exons using comprehensive human exon microarrays. Genome Biol. 8:R64.

Davis CA et al. 2018. The Encyclopedia of DNA elements (ENCODE): data portal update. Nucleic Acids Res. 46:D794–D801.

Dolphin AC. 2016. Voltage-gated calcium channels and their auxiliary subunits: physiology and pathophysiology and pharmacology. J. Physiol. 594:5369.

Dolphin AC. 2018. Voltage-gated calcium channel α 2δ subunits: an assessment of proposed novel roles. F1000Res. 7. doi: 10.12688/f1000research.16104.1.

Ewels P, Magnusson M, Lundin S, Käller M. 2016. MultiQC: summarize analysis results for multiple tools and samples in a single report. Bioinformatics. 32:3047–3048.

Forster JI et al. 2016. Characterization of Differentiated SH-SY5Y as Neuronal Screening Model Reveals Increased Oxidative Vulnerability. J. Biomol. Screen. 21:496–509.

Gallego-Paez LM et al. 2017. Alternative splicing: the pledge, the turn, and the prestige : The key role of alternative splicing in human biological systems. Hum. Genet. 136:1015–1042.

Gandal MJ et al. 2018. Transcriptome-wide isoform-level dysregulation in ASD, schizophrenia, and bipolar disorder. Science. 362. doi: 10.1126/science.aat8127.

Gao B et al. 2000. Functional properties of a new voltage-dependent calcium channel alpha(2)delta auxiliary subunit gene (CACNA2D2). J. Biol. Chem. 275:12237–12242.

Gleeson J, Lane TA, Harrison PJ, Haerty W, Clark MB. 2020. Nanopore direct RNA sequencing detects differential expression between human cell populations. Cold Spring Harbor Laboratory. 2020.08.02.232785. doi: 10.1101/2020.08.02.232785.

Grabowski P. 2011. Alternative splicing takes shape during neuronal development. Curr. Opin. Genet. Dev. 21:388–394.

Hardwick SA et al. 2016. Spliced synthetic genes as internal controls in RNA sequencing experiments. Nat. Methods. 13:792–798.

Hofmann F, Flockerzi V, Kahl S, Wegener JW. 2014. L-Type CaV1.2 Calcium Channels: From In Vitro Findings to In Vivo Function. Physiol. Rev. 94:303–326.

Jackson TC et al. 2020. Identification of Novel Targets of RBM5 in the Healthy and Injured Brain. Neuroscience. 440:299–315.

Jackson TC, Kochanek PM. 2020. RNA Binding Motif 5 (RBM5) in the CNS-Moving Beyond Cancer to Harness RNA Splicing to Mitigate the Consequences of Brain Injury. Front. Mol. Neurosci. 13:126.

Jaudon F et al. 2020. Targeting Alternative Splicing as a Potential Therapy for Episodic Ataxia Type 2. Biomedicines. 8. doi: 10.3390/biomedicines8090332.

Jiang L et al. 2011. Synthetic spike-in standards for RNA-seq experiments. Genome Res. 21:1543–1551.

Kelley LA, Mezulis S, Yates CM, Wass MN, Sternberg MJE. 2015. The Phyre2 web portal for protein modeling, prediction and analysis. Nat. Protoc. 10:845–858.

Kent WJ et al. 2002. The human genome browser at UCSC. Genome Res. 12:996–1006.

Keppetipola N, Sharma S, Li Q, Black DL. 2012. Neuronal regulation of pre-mRNA splicing by polypyrimidine tract binding proteins, PTBP1 and PTBP2. Crit. Rev. Biochem. Mol. Biol. 47:360–378.

Kim D, Langmead B, Salzberg SL. 2015. HISAT: a fast spliced aligner with low memory requirements. Nat. Methods. 12:357–360.

Kovalevich J, Langford D. 2013. Considerations for the use of SH-SY5Y neuroblastoma cells in neurobiology. Methods Mol. Biol. 1078:9–21.

Krueger F. 2015. Trim galore. https://www.bioinformatics.babraham.ac.uk/projects/trim_galore/.

Li H. 2018. Minimap2: pairwise alignment for nucleotide sequences. Bioinformatics. 34:3094–3100.

Linares AJ et al. 2015. The splicing regulator PTBP1 controls the activity of the transcription factor Pbx1 during neuronal differentiation. Elife. 4:e09268.

Liu J, Geng A, Wu X, Lin R-J, Lu Q. 2018. Alternative RNA Splicing Associated With Mammalian Neuronal Differentiation. Cereb. Cortex. 28:2810–2816.

McCarthy DJ, Chen Y, Smyth GK. 2012. Differential expression analysis of multifactor RNA-Seq experiments with respect to biological variation. Nucleic Acids Res. 40:4288–4297.

Mendsaikhan A, Takeuchi S, Walker DG, Tooyama I. 2018. Differences in Gene Expression Profiles and Phenotypes of Differentiated SH-SY5Y Neurons Stably Overexpressing Mitochondrial Ferritin. Front. Mol. Neurosci. 11:470.

Mészáros B, Erdos G, Dosztányi Z. 2018. IUPred2A: context-dependent prediction of protein disorder as a function of redox state and protein binding. Nucleic Acids Res. 46:W329–W337.

O’Bryan MK et al. 2013. RBM5 is a male germ cell splicing factor and is required for spermatid differentiation and male fertility. PLoS Genet. 9:e1003628.

Patro R, Duggal G, Love MI, Irizarry RA, Kingsford C. 2017. Salmon provides fast and bias-aware quantification of transcript expression. Nat. Methods. 14:417–419.

Punta M et al. 2012. The Pfam protein families database. Nucleic Acids Res. 40:D290–301.

Quinlan AR, Hall IM. 2010. BEDTools: a flexible suite of utilities for comparing genomic features. Bioinformatics. 26:841–842.

Raj B, Blencowe BJ. 2015. Alternative Splicing in the Mammalian Nervous System: Recent Insights into Mechanisms and Functional Roles. Neuron. 87:14–27.

R Core Team. 2016. R: A Language and Environment for Statistical Computing. Vienna, Austria https://www.R-project.org/.

Roy KR, Chanfreau GF. 2020. Robust mapping of polyadenylated and non-polyadenylated RNA 3’ ends at nucleotide resolution by 3’-end sequencing. Methods. 176:4–13.

Saito Y et al. 2019. Differential NOVA2-Mediated Splicing in Excitatory and Inhibitory Neurons Regulates Cortical Development and Cerebellar Function. Neuron. 101:707–720.e5.

Scotti MM, Swanson MS. 2016. RNA mis-splicing in disease. Nat. Rev. Genet. 17:19–32.

Sessegolo C et al. 2019. Transcriptome profiling of mouse samples using nanopore sequencing of cDNA and RNA molecules. Sci. Rep. 9:14908.

Shipley MM, Mangold CA, Szpara ML. 2016. Differentiation of the SH-SY5Y Human Neuroblastoma Cell Line. J. Vis. Exp. 53193.

Sibley CR, Blazquez L, Ule J. 2016. Lessons from non-canonical splicing. Nat. Rev. Genet. 17:407–421.

Soneson C et al. 2019. A comprehensive examination of Nanopore native RNA sequencing for characterization of complex transcriptomes. Nat. Commun. 10:3359.

Soneson C, Love MI, Robinson MD. 2015. Differential analyses for RNA-seq: transcript-level estimates improve gene-level inferences. F1000Res. 4:1521.

Splawski I et al. 2004. Ca(V)1.2 calcium channel dysfunction causes a multisystem disorder including arrhythmia and autism. Cell. 119:19–31.

Stark R, Grzelak M, Hadfield J. 2019. RNA sequencing: the teenage years. Nat. Rev. Genet. 20:631–656.

Sutherland LC, Wang K, Robinson AG. 2010. RBM5 as a putative tumor suppressor gene for lung cancer. J. Thorac. Oncol. 5:294–298.

Truckenmiller ME et al. 2001. Gene expression profile in early stage of retinoic acid-induced differentiation of human SH-SY5Y neuroblastoma cells. Restor. Neurol. Neurosci. 18:67–80.

Ule J et al. 2005. Nova regulates brain-specific splicing to shape the synapse. Nat. Genet. 37:844–852.

Van Nostrand EL et al. 2020. A large-scale binding and functional map of human RNA-binding proteins. Nature. 583:711–719.

Vitting-Seerup K, Sandelin A. 2019. IsoformSwitchAnalyzeR: analysis of changes in genome-wide patterns of alternative splicing and its functional consequences. Bioinformatics. 35:4469–4471.

Wang L et al. 2013. CPAT: Coding-Potential Assessment Tool using an alignment-free logistic regression model. Nucleic Acids Res. 41:e74.

Wang X et al. 2019. Full-length transcriptome reconstruction reveals a large diversity of RNA and protein isoforms in rat hippocampus. Nat. Commun. 10:5009.

Wang Z, Gerstein M, Snyder M. 2009. RNA-Seq: a revolutionary tool for transcriptomics. Nat. Rev. Genet. 10:57–63.

Warga RM, Wicklund A, Webster SE, Kane DA. 2016. Progressive loss of RacGAP1/ogre activity has sequential effects on cytokinesis and zebrafish development. Dev. Biol. 418:307–322.

Watanabe K, Taskesen E, van Bochoven A, Posthuma D. 2017. Functional mapping and annotation of genetic associations with FUMA. Nat. Commun. 8:1826.

Weyn-Vanhentenryck SM et al. 2018. Precise temporal regulation of alternative splicing during neural development. Nat. Commun. 9:2189.

Wong T, Deveson IW, Hardwick SA, Mercer TR. 2017. ANAQUIN: a software toolkit for the analysis of spike-in controls for next generation sequencing. Bioinformatics. 33:1723–1724.

Wyman D et al. 2020. A technology-agnostic long-read analysis pipeline for transcriptome discovery and quantification. bioRxiv. 672931. doi: 10.1101/672931.

Wyman D, Mortazavi A. 2019. TranscriptClean: variant-aware correction of indels, mismatches and splice junctions in long-read transcripts. Bioinformatics. 35:340–342.

Xu Y et al. 2019. Role of RNA-binding protein 5 in the diagnosis and chemotherapeutic response of lung cancer. Oncol. Lett. 17:2013–2019.

Yi L, Pimentel H, Bray NL, Pachter L. 2018. Gene-level differential analysis at transcript-level resolution. Genome Biol. 19:53.

Yuste R et al. 2020. A community-based transcriptomics classification and nomenclature of neocortical cell types. Nat. Neurosci. doi: 10.1038/s41593-020-0685-8.

Zhang Y-P et al. 2019. Down-regulated RBM5 inhibits bladder cancer cell apoptosis by initiating an miR-432-5p/β-catenin feedback loop. FASEB J. 33:10973–10985.

