## Supplementary Material for "Long read sequencing reveals novel isoforms and insights into splicing regulation during cell state changes"

**Supplementary Results & Discussion**

***Gene & transcript-level differential expression at more stringent log fold change thresholds***

Using more stringent threshold criteria (logFC $\gtreqless$ 1.5), 503 genes were upregulated in the differentiated cells, and 524 downregulated (Table S3), compared to the undifferentiated state (Fig 3A). These upregulated genes also showed greatest overlap with those up-regulated in the brain, compared with other tissue types ($p_{adj}$= 1.2 x $10^{-23}$). Whilst the permissive data showed significant overlap with genes down-regulated in brain, this relationship was not observed at the more stringent threshold ($p_{adj}$= 1).

At the transcript-level, we found a total of 884 transcripts significantly upregulated and 875 significantly downregulated in the differentiated cells (Fig 3C). These include 28 upregulated and 21 downregulated novel TALON transcripts respectively (Table S3).

**Supplementary Figures and Tables**

**Figure S1** Schematic representation of the custom annotation pipeline, utilising TALON software (Wyman et al. 2020) with custom bash, python and perl auxiliary and processing scripts (collated in clean_TALON_output.pl, see script repository).

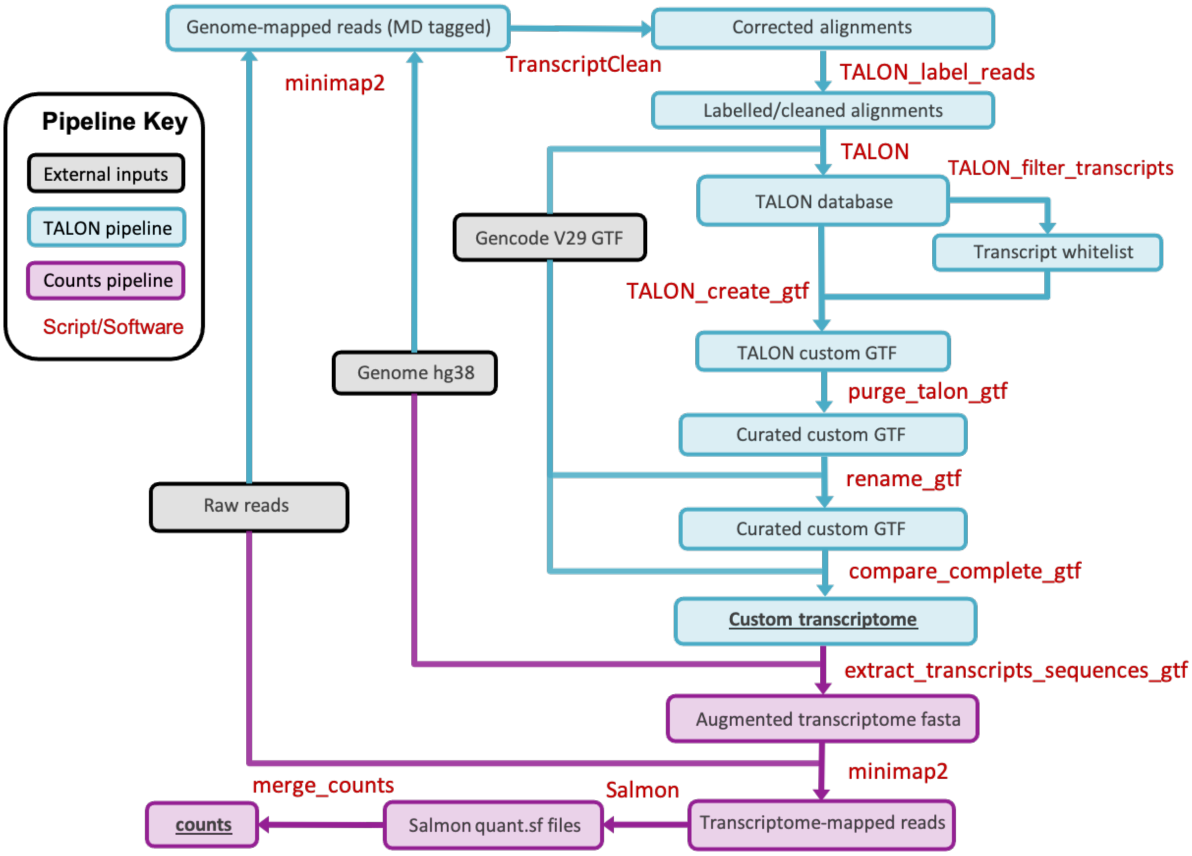

**Fig S2** Schematic representation of *CACNA2D2* (ENSG00000007402) transcripts, showing the novel transcript TALONT000703030. Figure modified from IsoformSwitchAnalyzeR output (Vitting-Seerup and Sandelin 2019).

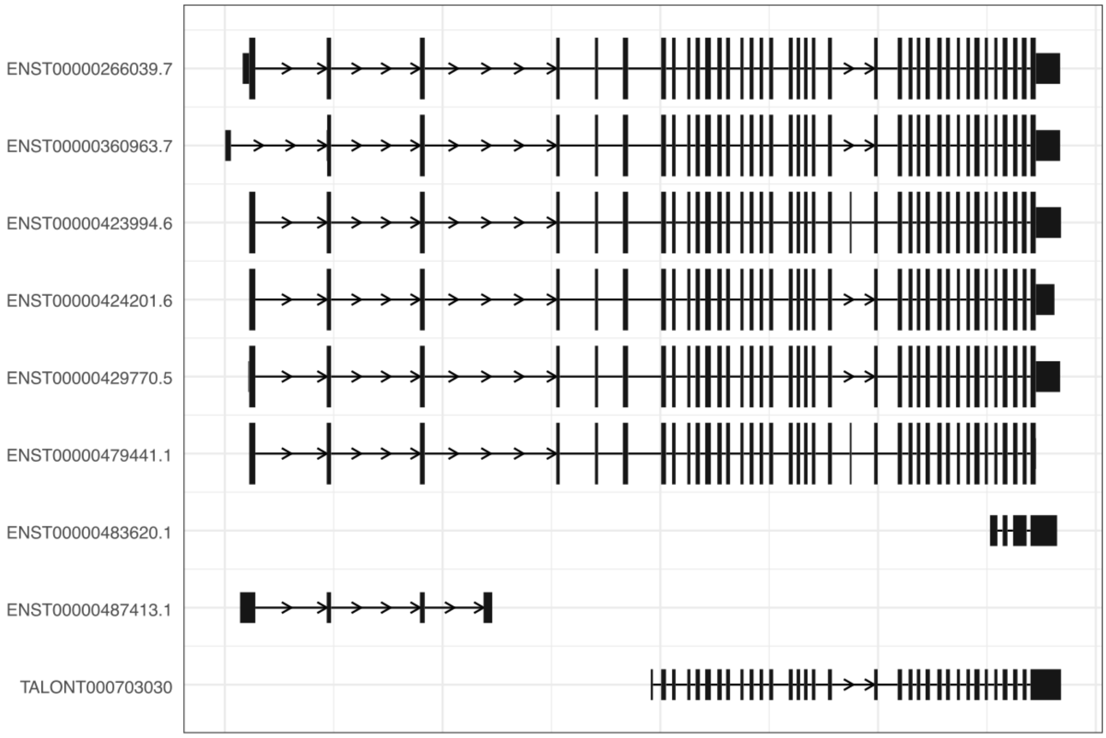


**Fig S3** Short read (Illumina paired-end) coverage plot of novel first exon (31bp) of CACNA2D2 (ENSG00000007402) transcript TALONT000703030 from all 10 sequencing runs (see Table S1).

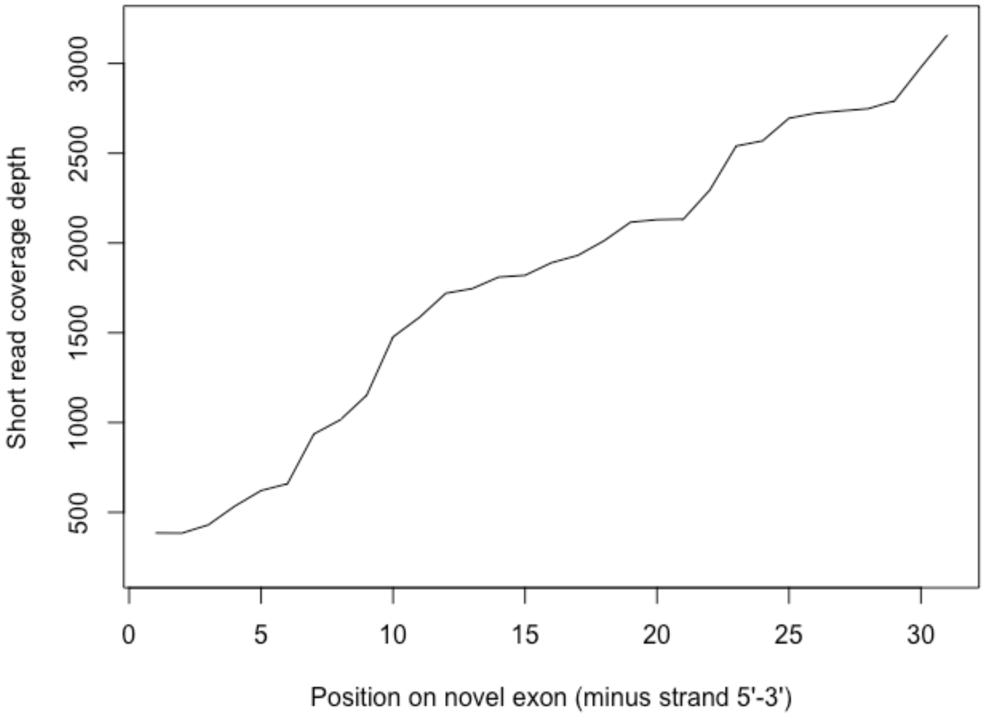

**Fig S4** Coverage plot of novel first exon (31bp) of CACNA2D2 (ENSG00000007402) transcript TALONT000703030 from N=27 human cortex RNA-seq GTEx accessions from N=21 individuals: SRR1310008, SRR1311400, SRR1311575, SRR1315866, SRR1316815, SRR1317344, SRR1320963, SRR1323043, SRR1326179, SRR1331579, SRR1333930, SRR1337564, SRR1339651, SRR1343481, SRR1353176, SRR1354446, SRR1364676, SRR1368772, SRR1382732, SRR1383059, SRR1387809, SRR1418837, SRR1418992, SRR1433971, SRR1435293, SRR1444580, SRR1468514.

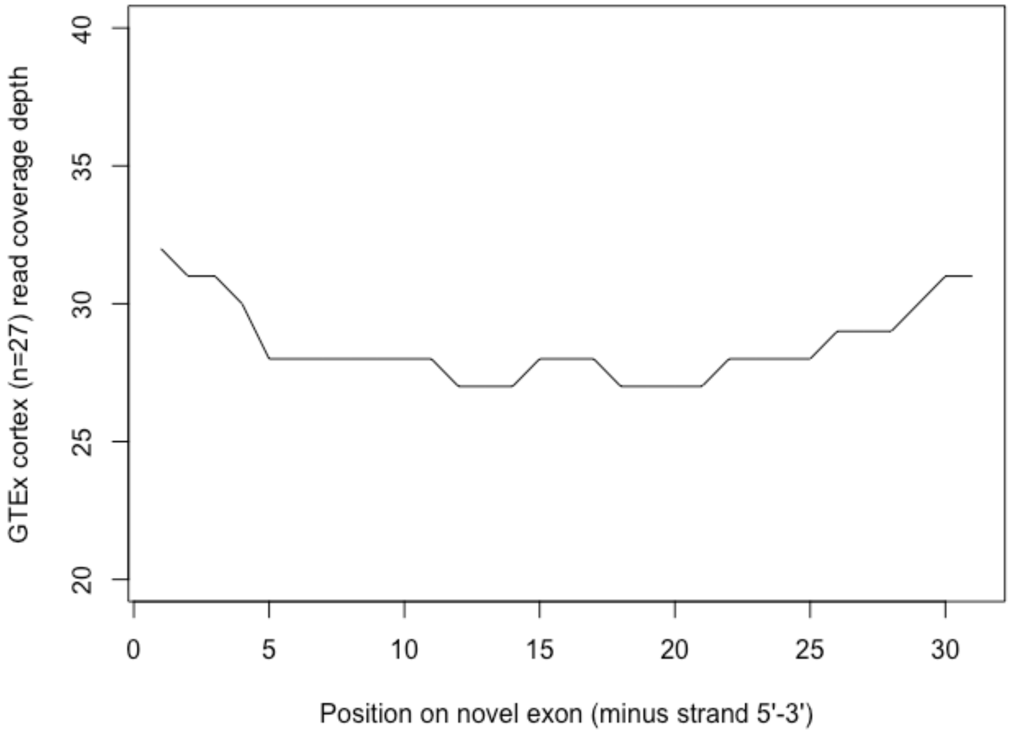


**Fig S5** Comparison of an example annotated coding transcript (ENST00000479441) of *CACNA2D2* (ENSG00000007402) with the novel transcript TALONT000703030, demonstrating key differences and initial 3D structure rendered using Phyre2 (Kelley et al. 2015).

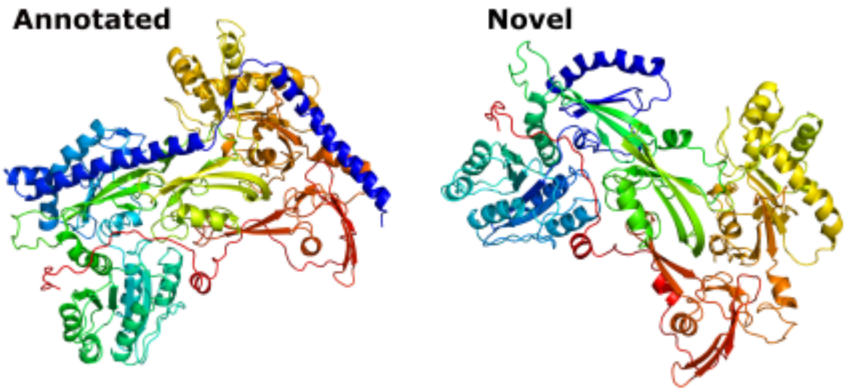

**Table S1** Summary statistics of the Oxford Nanopore Technologies (ONT) after quality checking and Illumina paired-end short read sequencing (SRS) of 10 replicate samples of human neuronal cell line SH-SY5Y. D = differentiated cell and U = undifferentiated cell samples.

| Sample ID | Reads | Bases | Median Length | Read N50 | Median Quality | Illumina Reads |
|---|---|---|---|---|---|---|
| D1 | 11,449,796 | 9,927,703,599 | 746 | 983 | 11.53 | 109,360,175 |
| D2 | 9,792,869 | 7,733,291,052 | 675 | 951 | 11.59 | 119,690,149 |
| D3 | 11,255,431 | 9,564,769,304 | 723 | 981 | 11.53 | 116,001,216 |
| D4 | 14,236,128 | 12,584,438,699 | 732 | 1016 | 11.59 | 61,118,341 |
| D5 | 11,503,329 | 9,379,344,639 | 707 | 961 | 11.64 | 98,946,695 |
| U1 | 9,171,036 | 7,737,779,710 | 701 | 1003 | 11.49 | 109,699,478 |
| U2 | 10,753,125 | 9,199,369,288 | 723 | 975 | 11.50 | 113,071,934 |
| U3 | 7,654,475 | 6,240,728,116 | 691 | 980 | 11.61 | 120,907,938 |
| U4 | 9,805,641 | 8,331,498,610 | 720 | 989 | 11.53 | 105,973,868 |
| U5 | 11,293,547 | 9,835,274,972 | 717 | 1023 | 11.54 | 98,721,393 |

**Table S2** Comparison of limit of quantification (LOQ) of Oxford Nanopore Technologies (ONT) sequencing, Illumina short read sequencing (SRS) and Illumina reads down-sampled to average nucleotide coverage of ONT reads. LOQ calculated by Anaquin (Wong et al. 2017).

| Sequencing Approach | LOQ genes (attoml/µl) | | LOQ isoforms (attoml/µl) | |
|---|---|---|---|---|
| | Sequin MixA | Sequin MixB | Sequin MixA | Sequin MixB |
| Illumina reads (**SRS**) | - | - | - | 0.0197 |
| Down-sampled Illumina | 0.118 | 0.118 | 0.059 | 0.0674 |
| Nanopore reads (**ONT**) | 0.118 | 0.472 | 0.059 | 0.270 |

**Table S3** Differential gene and transcript expression at a stringent filter of logFC ⋛ 1.5, FDR q-value < 0.05 (see also Figures 3A and 3C). Bracketed numbers refer to the portion of total that are TALON-identified novel transcripts. U = undifferentiated and D = differentiated cells, with arrows displaying expression directionality.

| Metric | Count | |
|---|---|---|
| | Gene level | Transcript level (Talon) |
| ↑U ↓D (≥ +1.5 logFC) | 524 | 875 (21) |
| ↑D ↓U (≤ -1.5 logFC) | 503 | 884 (28) |

**Table S4** N=104 Differential transcript usage switches with functional consequence ranked by q-value. Output from IsoformSwitchAnalyzeR (Vitting-Seerup and Sandelin 2019). See main methods for functional analysis.

| **Ensembl_geneID** | **gene_name** | **condition_1** | **condition_2** | **gene_switch_q_value** |
|---|---|---|---|---|
| ENSG00000086848 | ALG9 | Differentiated | Undifferentiated | 1.51E-26 |
| ENSG00000118363 | SPCS2 | Differentiated | Undifferentiated | 1.83E-21 |
| ENSG00000221838 | AP4M1 | Differentiated | Undifferentiated | 9.67E-18 |
| ENSG00000088986 | DYNLL1 | Differentiated | Undifferentiated | 4.99E-12 |
| ENSG00000167113 | COQ4 | Differentiated | Undifferentiated | 1.25E-11 |
| ENSG00000112769 | LAMA4 | Differentiated | Undifferentiated | 1.41E-11 |
| ENSG00000164117 | FBXO8 | Differentiated | Undifferentiated | 1.95E-10 |
| ENSG00000101460 | MAP1LC3A | Differentiated | Undifferentiated | 1.57E-08 |
| ENSG00000108384 | RAD51C | Differentiated | Undifferentiated | 4.19E-08 |
| ENSG00000172053 | QARS | Differentiated | Undifferentiated | 1.08E-07 |
| ENSG00000169738 | DCXR | Differentiated | Undifferentiated | 2.91E-07 |
| ENSG00000160789 | LMNA | Differentiated | Undifferentiated | 3.67E-07 |
| ENSG00000140416 | TPM1 | Differentiated | Undifferentiated | 7.53E-07 |
| ENSG00000172081 | MOB3A | Differentiated | Undifferentiated | 1.23E-06 |

| | | | | |
|---|---|---|---|---|
| ENSG00000151148 | UBE3B | Differentiated | Undifferentiated | 1.52E-06 |
| ENSG00000146282 | RARS2 | Differentiated | Undifferentiated | 2.19E-06 |
| ENSG00000221926 | TRIM16 | Differentiated | Undifferentiated | 6.06E-06 |
| ENSG00000163738 | MTHFD2L | Differentiated | Undifferentiated | 2.28E-05 |
| ENSG00000108591 | DRG2 | Differentiated | Undifferentiated | 3.53E-05 |
| ENSG00000179632 | MAF1 | Differentiated | Undifferentiated | 3.79E-05 |
| ENSG00000158604 | TMED4 | Differentiated | Undifferentiated | 4.45E-05 |
| ENSG00000068024 | HDAC4 | Differentiated | Undifferentiated | 9.36E-05 |
| ENSG00000285437 | POLR2J3 | Differentiated | Undifferentiated | 0.000105381 |
| ENSG00000125755 | SYMPK | Differentiated | Undifferentiated | 0.000216866 |
| ENSG00000056972 | TRAF3IP2 | Differentiated | Undifferentiated | 0.000341384 |
| ENSG00000089486 | CDIP1 | Differentiated | Undifferentiated | 0.000365387 |
| ENSG00000163053 | SLC16A14 | Differentiated | Undifferentiated | 0.000516673 |
| ENSG00000171469 | ZNF561 | Differentiated | Undifferentiated | 0.000538662 |
| ENSG00000055044 | NOP58 | Differentiated | Undifferentiated | 0.000654218 |
| ENSG00000107317 | PTGDS | Differentiated | Undifferentiated | 0.000728674 |
| ENSG00000165792 | METTL17 | Differentiated | Undifferentiated | 0.000728674 |
| ENSG00000127125 | PPCS | Differentiated | Undifferentiated | 0.00074317 |
| ENSG00000184347 | SLIT3 | Differentiated | Undifferentiated | 0.000875632 |
| ENSG00000104870 | FCGRT | Differentiated | Undifferentiated | 0.000913016 |
| ENSG00000180596 | HIST1H2BC | Differentiated | Undifferentiated | 0.000978953 |
| ENSG00000131711 | MAP1B | Differentiated | Undifferentiated | 0.0010522 |
| ENSG00000179029 | TMEM107 | Differentiated | Undifferentiated | 0.0010522 |
| ENSG00000140995 | DEF8 | Differentiated | Undifferentiated | 0.001148515 |
| ENSG00000135931 | ARMC9 | Differentiated | Undifferentiated | 0.001238557 |

| | | | | |
|---|---|---|---|---|
| ENSG00000161800 | RACGAP1 | Differentiated | Undifferentiated | 0.001494161 |
| ENSG00000146540 | C7orf50 | Differentiated | Undifferentiated | 0.001536413 |
| ENSG00000183092 | BEGAIN | Differentiated | Undifferentiated | 0.001547641 |
| ENSG00000123178 | SPRYD7 | Differentiated | Undifferentiated | 0.001966476 |
| ENSG00000003756 | RBM5 | Differentiated | Undifferentiated | 0.00235666 |
| ENSG00000166444 | ST5 | Differentiated | Undifferentiated | 0.00235666 |
| ENSG00000124222 | STX16 | Differentiated | Undifferentiated | 0.002830477 |
| ENSG00000196704 | AMZ2 | Differentiated | Undifferentiated | 0.002854375 |
| ENSG00000164466 | SFXN1 | Differentiated | Undifferentiated | 0.00306238 |
| ENSG00000198408 | OGA | Differentiated | Undifferentiated | 0.003637501 |
| ENSG00000197372 | ZNF675 | Differentiated | Undifferentiated | 0.004074968 |
| ENSG00000099330 | OCEL1 | Differentiated | Undifferentiated | 0.004090568 |
| ENSG00000257591 | ZNF625 | Differentiated | Undifferentiated | 0.00422435 |
| ENSG00000064490 | RFXANK | Differentiated | Undifferentiated | 0.004622242 |
| ENSG00000136490 | LIMD2 | Differentiated | Undifferentiated | 0.006030149 |
| ENSG00000131477 | RAMP2 | Differentiated | Undifferentiated | 0.006540547 |
| ENSG00000184787 | UBE2G2 | Differentiated | Undifferentiated | 0.006598612 |
| ENSG00000196465 | MYL6B | Differentiated | Undifferentiated | 0.007142517 |
| ENSG00000132692 | BCAN | Differentiated | Undifferentiated | 0.007231216 |
| ENSG00000138606 | SHF | Differentiated | Undifferentiated | 0.007762126 |
| ENSG00000151353 | TMEM18 | Differentiated | Undifferentiated | 0.007989728 |
| ENSG00000197122 | SRC | Differentiated | Undifferentiated | 0.008177219 |
| ENSG00000160013 | PTGIR | Differentiated | Undifferentiated | 0.008905808 |
| ENSG00000111581 | NUP107 | Differentiated | Undifferentiated | 0.009183614 |
| ENSG00000197771 | MCMBP | Differentiated | Undifferentiated | 0.009700325 |

| | | | | |
|---|---|---|---|---|
| ENSG00000137434 | C6orf52 | Differentiated | Undifferentiated | 0.009711457 |
| ENSG00000198551 | ZNF627 | Differentiated | Undifferentiated | 0.00989504 |
| ENSG00000066379 | ZNRD1 | Differentiated | Undifferentiated | 0.01087531 |
| ENSG00000062194 | GPBP1 | Differentiated | Undifferentiated | 0.011249117 |
| ENSG00000166471 | TMEM41B | Differentiated | Undifferentiated | 0.011305356 |
| ENSG00000174628 | IQCK | Differentiated | Undifferentiated | 0.012469732 |
| ENSG00000227051 | C14orf132 | Differentiated | Undifferentiated | 0.013453422 |
| ENSG00000141298 | SSH2 | Differentiated | Undifferentiated | 0.013885209 |
| ENSG00000121413 | ZSCAN18 | Differentiated | Undifferentiated | 0.014960611 |
| ENSG00000151617 | EDNRA | Differentiated | Undifferentiated | 0.016912634 |
| ENSG00000166407 | LMO1 | Differentiated | Undifferentiated | 0.017290737 |
| ENSG00000100888 | CHD8 | Differentiated | Undifferentiated | 0.01754742 |
| ENSG00000276234 | TADA2A | Differentiated | Undifferentiated | 0.018780264 |
| ENSG00000198663 | C6orf89 | Differentiated | Undifferentiated | 0.020507021 |
| ENSG00000205339 | IPO7 | Differentiated | Undifferentiated | 0.020514719 |
| ENSG00000136146 | MED4 | Differentiated | Undifferentiated | 0.02127913 |
| ENSG00000004897 | CDC27 | Differentiated | Undifferentiated | 0.022166596 |
| ENSG00000119559 | C19orf25 | Differentiated | Undifferentiated | 0.024137326 |
| ENSG00000060982 | BCAT1 | Differentiated | Undifferentiated | 0.024277036 |
| ENSG00000196482 | ESRRG | Differentiated | Undifferentiated | 0.027583627 |
| ENSG00000139182 | CLSTN3 | Differentiated | Undifferentiated | 0.027739353 |
| ENSG00000204653 | ASPDH | Differentiated | Undifferentiated | 0.028996139 |
| ENSG00000106635 | BCL7B | Differentiated | Undifferentiated | 0.029720469 |
| ENSG00000144401 | METTL21A | Differentiated | Undifferentiated | 0.029949427 |
| ENSG00000135090 | TAOK3 | Differentiated | Undifferentiated | 0.033871798 |

| | | | | |
|---|---|---|---|---|
| ENSG00000149292 | TTC12 | Differentiated | Undifferentiated | 0.033913158 |
| ENSG00000171953 | ATPAF2 | Differentiated | Undifferentiated | 0.034218455 |
| ENSG00000080986 | NDC80 | Differentiated | Undifferentiated | 0.034297085 |
| ENSG00000105321 | CCDC9 | Differentiated | Undifferentiated | 0.034903122 |
| ENSG00000135541 | AHI1 | Differentiated | Undifferentiated | 0.035365825 |
| ENSG00000122490 | PQLC1 | Differentiated | Undifferentiated | 0.036240151 |
| ENSG00000174353 | STAG3L3 | Differentiated | Undifferentiated | 0.038682053 |
| ENSG00000140939 | NOL3 | Differentiated | Undifferentiated | 0.04412664 |
| ENSG00000147548 | NSD3 | Differentiated | Undifferentiated | 0.044278588 |
| ENSG00000104231 | ZFAND1 | Differentiated | Undifferentiated | 0.044308667 |
| ENSG00000107951 | MTPAP | Differentiated | Undifferentiated | 0.044598461 |
| ENSG00000166398 | KIAA0355 | Differentiated | Undifferentiated | 0.046626246 |
| ENSG00000259431 | THTPA | Differentiated | Undifferentiated | 0.047528021 |
| ENSG00000135249 | RINT1 | Differentiated | Undifferentiated | 0.047949428 |
| ENSG00000136574 | GATA4 | Differentiated | Undifferentiated | 0.048524879 |

### CITATIONS


Kelley, Lawrence A., Stefans Mezulis, Christopher M. Yates, Mark N. Wass, and Michael J. E. Sternberg. 2015. “The Phyre2 Web Portal for Protein Modeling, Prediction and Analysis.” *Nature Protocols* 10 (6): 845–58.

Vitting-Seerup, Kristoffer, and Albin Sandelin. 2019. “IsoformSwitchAnalyzeR: Analysis of Changes in Genome-Wide Patterns of Alternative Splicing and Its Functional Consequences.” *Bioinformatics* 35 (21): 4469–71.

Wong, Ted, Ira W. Deveson, Simon A. Hardwick, and Tim R. Mercer. 2017. “ANAQUIN: A Software Toolkit for the Analysis of Spike-in Controls for next Generation Sequencing.” *Bioinformatics* 33 (11): 1723–24.

Wyman, Dana, Gabriela Balderrama-Gutierrez, Fairlie Reese, Shan Jiang, Sorena Rahmanian, Stefania Forner, Dina Matheos, et al. 2020. “A Technology-Agnostic Long-Read Analysis Pipeline for Transcriptome Discovery and Quantification.” *bioRxiv*. https://doi.org/10.1101/672931.
